# Phosphoproteomics uncovers rapid and specific transition from plant two-component system signaling to Ser/Thr phosphorylation by the intracellular redox sensor AHK5

**DOI:** 10.1101/2025.10.13.682113

**Authors:** Thomas Drechsler, Zhi Li, Waltraud X. Schulze, Klaus Harter

## Abstract

The *Arabidopsis thaliana* histidine kinase 5 (AHK5) is proposed to function as a intracellular sensor of active oxygen species (ROS) and redox conditions, fine-tuning responses to diverse ROS-inducing stimuli through a multistep phosphorelay (MSP) system. To elucidate the mechanism of ROS perception by AHK5 and signaling through posttranslational modifications, a comparative phosphoproteomics analysis was conducted using *ahk5* loss-of-function and wild type seedlings in response to exogenously applied H_2_O_2_. Under control and H_2_O_2_-related conditions, a rapid signaling transition from the MSP to Ser/Thr phosphorylation was observed. AHK5 appears to regulate ROS-responsive phosphorylation at the plasma membrane by the modification of nanodomain components such as remorins, aquaporins, IQDs, and patellins, positioning it as a central integrator of redox signaling during adaptation to diverse stresses. AHK5 drives a distinct, ABA-independent signaling pathway for H_2_O_2_-induced stomatal closure, likely by regulating phosphorylation of key components such as RBOHD, CAS, and HPCA1. Moreover, AHK5-fine-tuned root development involves ABA-dependent signaling and appears to act through modulation of auxin transport *via* phosphorylation of auxin carrier, highlighting a mechanistic divergence between ROS responses in the shoot and root. Our data further suggest that AHK5 orchestrates stress-responsive nanodomain-associated signaling hubs through regulation of membrane trafficking, cytoskeletal dynamics, and endocytosis to mediate stress responses.

**Highlight:** AHK5 perceives and may amplify H_2_O_2_ signaling by linking two-component systems to Ser/Thr phosphorylation, likely by modulating membrane-associated signalosomes, thereby fine-tuning responses to stress-induced reactive oxygen species.

## Introduction

As byproducts of the Photosynthetic Electron Transport Chain (PETC), Respiratory Electron Transport Chain (RETC), and photorespiration, Reactive Oxygen Species (ROS) are recognized as key second messengers of intra- and intercellular signaling (Waszczak *et al*., 2018; Huang *et al*., 2019; Smirnoff and Arnaud, 2019; Choudhary *et al*., 2020; Huang *et al*., 2020; Lee *et al*., 2020; Sies and Jones, 2020; Meyer *et al*., 2021; Wang *et al*., 2021; Marcec and Tanaka, 2022). Among these, comparably long-lived hydrogen peroxide (H_2_O_2_), has become widely accepted as key integrator of external and internal cues. The most abundant ROS in plants, the superoxide anion (O_2_^-^), is predominantly generated by photoreduction of oxygen in thylakoids during photosynthesis (Asada, 2006). During stress responses and development, especially O_2_^-^ and H₂O₂ act as the main agents in maintaining cellular redox homeostasis in plants in conjunction with the glutathione redox buffer system (for reviews see He *et al*. (2018); Waszczak *et al*. (2018); Huang *et al*. (2020)).

One of the first responses to biotic stress, such as pathogen attack, is the generation of high amounts of ROS, the so-called ROS burst. In the apoplast, plasma membrane (PM) anchored calcium-dependent NADPH oxidases, the RESPIRATORY BURST OXIDASE HOMOLOGS (RBOH) increase ROS levels to combat infection by damaging oxidation of bacterial and fungal cell wall components (Doke, 1983; Apostol *et al*., 1989; Levine *et al*., 1994). RBOHs release O_2_^-^ into the apoplast, where it is subsequently either spontaneously or enzymatically dismutated into H₂O₂, which can diffuse back into the cell through aquaporins, altering intracellular redox characteristics and initiating intracellular signaling (Desikan *et al*., 1996; Asada, 2006; Bienert *et al*., 2006; Bienert *et al*., 2007; Dynowski *et al*., 2008). Other processes utilizing ROS, especially O_2_^-^ and H_2_O_2_, as second messengers are abiotic stress responses, root development, including root hair and lateral root formation, organ senescence, and stomatal closure (Zhang *et al*., 2001; Neill *et al*., 2002; Kwak *et al*., 2003; Desikan *et al*., 2006; Zimmermann *et al*., 2006; Verslues *et al*., 2007; Neill *et al*., 2008; Clark *et al*., 2010; Hua *et al*., 2012; Zou *et al*., 2015; Heunemann, 2016; Wu *et al*., 2020; Fichman *et al*., 2022). The plethora of processes regulated by ROS highlights the importance of accurate mechanisms to sense and adjust ROS levels during response to environmental and internal cues.

Plants feature intracellular redox buffers to maintain redox homeostasis, which include non-enzymatic flavonoids or reduced/oxidated glutathione (GSH/GSSG) and ascorbate/dehydroascorbate (ASC/DHA) redox couples, as well as enzymatic ascorbate peroxidases (APX), thioredoxins (TRX), glutaredoxins (GRX), and catalases (CAT). These systems contribute to specific intracellular redox characteristics in response to diverse stimuli, including those overloading the redox buffer system resulting in ‘oxidative stress’ (for revies see Kapoor *et al*. (2015); He *et al*. (2018); Waszczak *et al*. (2018)). Recently, a membrane anchored ROS sensor with an extracellular ROS sensing domain, the Leucine-Rich Repeat (LRR) Receptor Like Kinase (RLK) HYDROGEN PEROXIDE INDUCED CALCIUM INCREASES 1 (HPCA1), was discovered. HPCA1 gets oxidized upon external hydrogen (eH_2_O_2_) perception at extracellular cysteine residues, leading to its dimerization and subsequent activation of kinase activity, in turn activating Ca^2+^ influx channels (Wu *et al*., 2020). HPCA1 may be required for intercellular and systemic propagation of Ca^2+^ and apoplastic H_2_O_2_ signals (Fichman *et al*., 2022). Emerging evidence suggests, that characteristic ROS signatures, varying in amplitude and frequency, may encode some degree of specificity that leads to distinct responses (Mittler *et al*., 2011; Gilroy *et al*., 2014; Wang *et al*., 2021).

Phosphorylation is one of the main modes of altering protein function in living organisms, also in the context of stress signaling. An evolutionary old mechanism of signaling *via* protein phosphorylation cascades is the two-component system (TCS). The basic TCS, ubiquitously present in prokaryotes, consists of two components: A Histidine Kinase (HK) that autophosphorylates at a conserved histidine (His) residue upon signal perception and a Response Regulator (RR) with a conserved aspartate (Asp) to which the phosphoryl group is transferred to. The RR in turn modulates responses either by acting as a transcription factor or through changes in protein-protein interactions (Mira-Rodado, 2019). The TCS in plants has evolved into a more refined version, employing additional phosphorylation steps, called the multistep phosphorelay (MSP) system. In plant MSP, there is an initial intramolecular transfer of the phosphate from His(p) of the HK kinase domain of the HK to a conserved Asp in the receiver domain within the same molecule. Hence, they are called Hybrid Histidine Kinases (HHKs). Before transfer of the phosphoryl group from Asp(p) to a RR, the phosphate is at first relayed to a conserved histidine of histidine-containing phosphotransfer proteins (HPs) from which it is again transferred to the Asp of the RR (Mira-Rodado, 2019). Importantly, a direct transfer of the phosphoryl residue from N-phosphoamidates like His or Asp to O-phosphomonoesters like Ser, Thr, or Tyr residues and *vice versa* is not possible (Dautel *et al*., 2016). Therefore, Ser/Thr/Tyr phosphorylation caused by MSP elements must occur via induction of classical kinase activity probably by protein-protein interaction (Dautel *et al.,* 2016).

The MSP of *Arabidopsis* consists of 11 ARABIDOPSIS HISTIDINE KINASES (AHKs), 6 ARABIDOPSIS HISTIDINE-CONTAINING PHOSPHOTRANSFER PROTEINS (AHPs), and 23 ARABIDOPSIS RESPONSE REGULATORES (ARRs) (Mira-Rodado, 2019). *Arabidopsis* HKs and HK-like proteins can be divided into 3 main groups: Ethylene receptors, phytochromes, and AHKs. AHK2 to 4 act as primary cytokinin receptors (Yamada *et al*., 2001; Higuchi *et al*., 2004), whereas AHK1 plays a role during osmotic stress responses, possibly acting as a mechanosensor, and also appears to be involved in brassinosteroid signaling (Tran *et al*., 2007; Dautel, 2016). In contrast to the other membrane anchored AHKs, AHK5 does not feature intermembrane loops and an extracellular domain, resulting in a nucleo-cytoplasmic localization (Desikan *et al*., 2008; Heunemann, 2016). AHK5 participates in various ROS-related responses such as leaf senescence, root growth, defense to pathogen attack, and stomatal closure either directly in response to H_2_O_2_- or ROS-inducing stimuli like flg22, ET, darkness, salt stress, or pathogen infection (Iwama *et al*., 2007; Desikan *et al*., 2008; Mira-Rodado *et al*., 2012; Pham and Desikan, 2012; Pham *et al*., 2012; Heunemann, 2016). Since no altered H_2_O_2_ levels were observed in an AHK5 loss-of-function mutant (*ahk5-1)* under non-stress conditions, it was proposed that *ahk5-1* plants are defective in proper H_2_O_2_ sensing and that AHK5 may act as an intracellular ROS sensor (Heunemann, 2016). The proposed redox-sensing function is further supported by the presence of a redox-sensitive cysteine at position 3 (C3) of the amino acid sequence, which enables the *in vitro* dimerization of the AHK5 input domain to various degrees corresponding with different concentrations of H_2_O_2_ and GSH/GSSG redox potentials (Heunemann, 2016; Drechsler, 2022; Nöldeke, 2022).

For the PM-located AHK1, the signaling transition from the MSP with its energetically unstable His/Asp phosphorylation to more energetically stable Ser/Thr/Tyr phosphorylation cascades was shown recently (Dautel, 2016). Similar signaling transitions were also demonstrated for the cytokinin receptors AHK2 and AHK3 (Dautel *et al*., 2016). Here we show that also AHK5 proceeds massively into multiple Ser/Thr/Tyr phosphorylation pathways. Thereby, it likely modifies the nanoscale organization of PM proteins as well as the activity and integrity of several signaling networks required for appropriate H_2_O_2_- and redox potential-dependent regulation of abiotic and biotic stress responses

## Materials and methods

*Arabidopsis thaliana* lines *ahk5-1* (Col-0) seeds were kindly provided by Dr. Radhika Desikan (Desikan *et al*., 2008). ^15^N and ^14^N potassium nitrate as the metabolic labeling supplement was ordered from Cambridge Isotope Laboratories, Inc. The procedure for hydroponic growth of *A. thaliana* seedlings was adapted from (Kierszniowska *et al*., 2008; Wu and Schulze, 2015; Dautel, 2016; Dautel *et al*., 2016; Fischer, 2018). In brief, 20 mg per sample of surface sterilized seeds using chloric gas were stratified in JPL medium containing the appropriate ^14^N/^15^N nitrogen source at 4 °C in darkness for 3 days and transferred to 250 ml Erlenmeyer flasks containing 50 ml of JPL growth medium. Wild type (Col-0) seedlings were grown with the addition of ^14^N as the only nitrogen source, whereas *ahk5-1* seedlings were supplied with ^15^N. Flasks were transferred to 22 °C constant light on a shaker (80 rpm). After 10 days after germination (dag), growth medium was exchanged with fresh JPL and the seedlings grown for an additional 4 days before treatment (14 dag total). For the treatment, always starting at 1 pm mid-day, growth medium was exchanged with fresh JPL containing 5 mM freshly added H_2_O_2_ (“eH_2_O_2_”) or the corresponding volume of water (“mock”) and incubated for 10 min shaking at 80 rpm constant light before immediately rinsing with tap water, gently drying on paper towels and deep freezing in liquid nitrogen. Samples were kept at -80 °C until protein extraction.

Protein extraction, tryptic digestion (in-solution) and phosphopeptide enrichment was executed according to a published procedure (Wu and Schulze, 2015) with the following modifications: Instead of grinding in liquid nitrogen with mortar and pestle, frozen plant material was coarsely crushed and directly subjected to manual potter tissue homogenization in 10 ml microsome isolation buffer and thoroughly homogenized while avoiding air bubbles. Samples were transferred to 50 ml falcon tubes and centrifuged for 15 min at 4 °C at 7500 g. Supernatant was filtered through miracloth and subjected to ultracentrifugation for microsomal pelleting. Thereafter, sample processing was continued according to the protocol (Wu and Schulze, 2015; Dautel, 2016).

MS analysis and subsequent calculation of differentially phosphorylated phosphopeptides was carried out as described before (Wu and Schulze, 2015; Dautel, 2016) for the metabolic labeling experiment with the following modifications: Acquired spectra were matched against the *Arabidopsis* proteome (TAIR10, 35386 entries) using MaxQuant v.1.4.1.2. Carbamidomethylation of cysteine was set as fixed modification and oxidation of methionine as well as phosphorylation of serine, threonine and tyrosine was set as variable modifications. Mass tolerance for the database search was set to 20 ppm on full scans and 0.5 Da for fragment ions. Peptides were accepted using an FDR threshold of 0.01. The option “matched features” was set, and these features were extracted and matched to identified ^14^N-form spectra as described in Pertl-Obermeyer *et al*. (2016). For quantitation, ratios between heavy (^15^N) and light (^14^N) forms of each peptide were calculated using an in-house script (Pertl-Obermeyer *et al*., 2016). Hits to contaminants (e.g. keratins) and additionally identified reverse hits were excluded from further analysis. Ratios indicate “mutant vs wild type” ratios and were averaged. All phosphopeptides and proteotypic non-phosphopeptides were used for quantitation. Within each sample, log_2_(^15^N/^14^N) values were calculated for each peptide ions species (each m/z) and these ratios were averaged among at least three biological replicates. All phosphopeptides derived from 35S::GFP-AHK5 plants were used for quantitation, but not considered for further analysis, due to posttranscriptional silencing. The distribution of log_2_ ratios was normalized to be centered on 0. Due to ^14^N/^15^N metabolic labeling, the entire sample processing and MS analysis could be carried out in combined samples of material of both *ahk5-1* and WT genotypes. Hence, a direct comparison of log_2_(^15^N/^14^N) ratios (log_2_FC) was possible without calculating p-values, in which positive values indicate more phosphorylation and negative values indicate less phosphorylation in *ahk5-1* compared to WT. Since there was no parallel experiment using reciprocal metabolic labeling, log_2_FC values above 2.0 or below -2.0 were considered as phosphopeptides differentially phosphorylated between wild type and mutants. Further, phosphopeptides were considered responsive to treatment (“eH_2_O_2_ dependent”), if the difference of log_2_FC values between mock and eH_2_O_2_ treated samples was at least 1, translating into at least two-fold increase or decrease in phosphorylation levels between the two conditions. Phosphopeptides were considered unresponsive to treatment (“eH_2_O_2_ independent”), if the log_2_FC difference was 1 or less.

## Results and Discussion

### AHK5 causes rapid Ser/Thr phosphorylation in response to eH_2_O_2_

Given AHK5’s established role in stress signaling, particularly in H_2_O_2_-mediated pathways (Iwama *et al*., 2007; Desikan *et al*., 2008; Pham and Desikan, 2012; Pham *et al*., 2012; Heunemann, 2016), both hydroponically grown and differentially metabolically labeled *ahk5-1* (^15^N) and wild-type (^14^N) seedlings were treated with 5 mM eH_2_O_2_ or mock treated for 10 minutes in parallel experiments before harvest. Due to metabolic labeling, downstream sample processing could be carried out in combined samples of the different genotypes, which allows for a direct quantitative comparison of the (phospho-) proteome of *ahk5-1* and wild-type (WT) seedlings. After protein extraction and phosphopeptide enrichment, quantitative phosphoproteomic analysis identified several differentially phosphorylated phosphopeptides. The histogram of normalized log_2_FC values centered on zero indicates, that the majority of phosphopeptides were not affected by the absence of AHK5 or the eH_2_O_2_ treatment. Phosphopeptides with a log_2_FC above 2.0 or below -2.0, which was outside the width of the standard deviation (Fig. 1A) were considered as phosphopeptides differentially phosphorylated between the *ahk5-1* mutant and WT (Fig. 1B). Further, phosphopeptides were considered responsive to treatment, if (“eH_2_O_2_ dependent”), if the log_2_FC values of the *ahk5-1* mutant versus WT in mock and eH_2_O_2_ treated samples were at least two-fold different (Fig. 1B, cyan dots).

**Fig. 1:**
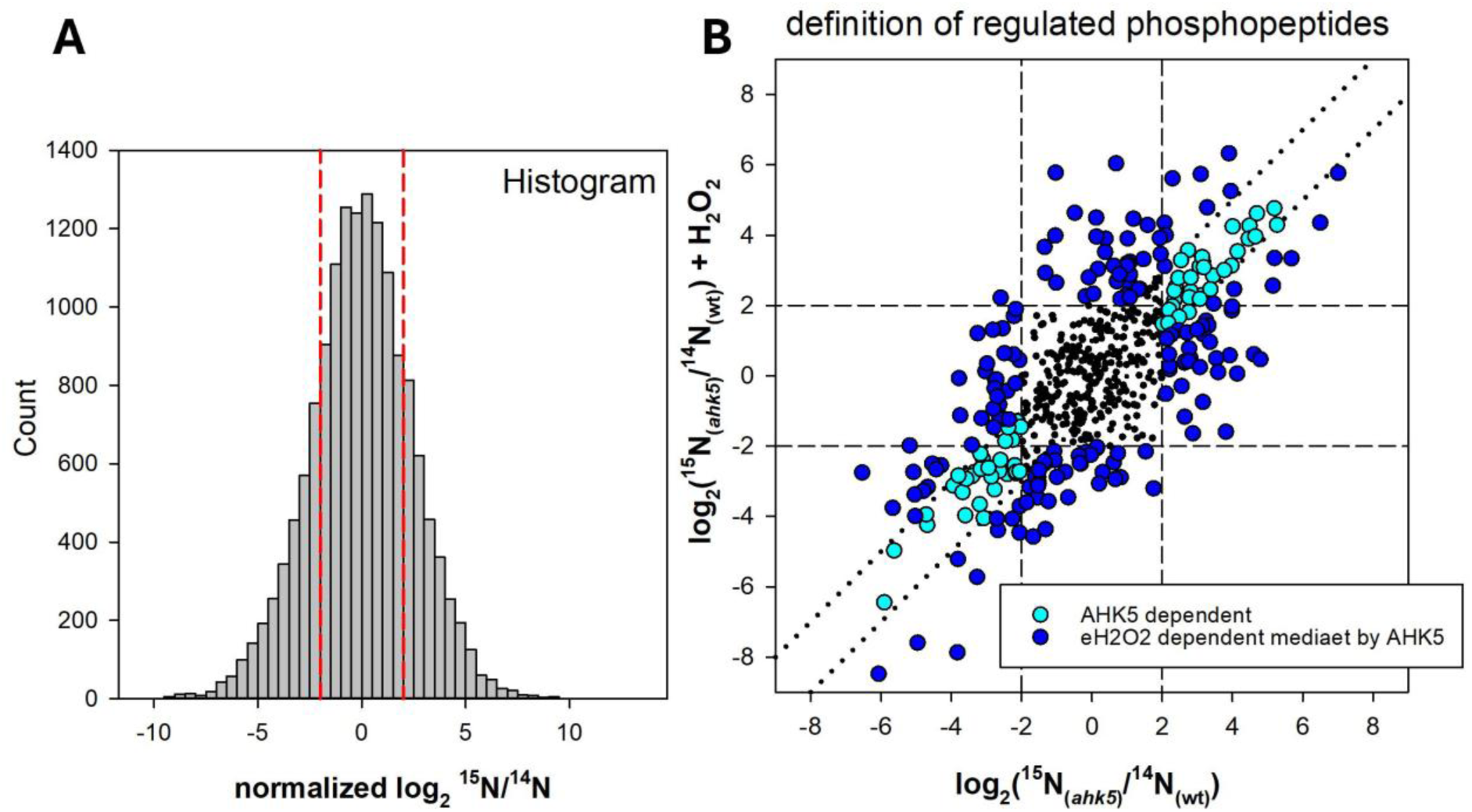
Definition of AHK5- and H2O2-dependent phopsphopeptides. (**A**) Histogram of normalized log2(^15^N/^14^N) ratios in which ^15^N represents *ahk5-1* proteins and ^14^N represents WT proteins. The red lines indicate the width of one standard deviation, which was used as a cutoff-threshold. (**B**) Cross plot of log2 ratios of phosphopeptide intensities of mutant (*ahk5-1*) vs. WT in mock-treated and eH2O2 treated experiments as derived from respective ^15^N/^14^N ratios. Each dot represents one phosphopeptide. Cyan: phosphopeptides consistently altered in mutant vs. WT independent of treatment. Blue: phosphopeptides differentially phosphorylated in mutant vs. WT AND with log2 fold change mutant vs WT least factor 2 different in H2O2 treatment compared to control.

A total of 1,159 phosphopeptides were quantified across all analyzed samples (Fig. 2). Of these, 1,020 contained at least one phosphoserine, 153 contained at least one phosphothreonine, and a single peptide featured a phosphotyrosine residue. Phosphopeptides were classified as differentially phosphorylated (DPP) based on an absolute log₂ fold change (FC) greater than 2 (Dautel, 2016; Dautel *et al*., 2016). Using this criterion, 266 DPPs were identified in mock treated *ahk5-1* plants compared to WT plants, while 279 DPPs were identified in *ahk5-1* plants compared to wild-type plants in response to eH_2_O_2_. Functional categorization of the proteins corresponding to these phosphopeptides was performed using MapMan. Disregarding the category “other”, the categories “protein.posttranslational modification”, “RNA.regulation of transcription”, “signaling.receptor kinases”, and “transport” shared more than 50% of DPs (Fig. 2).

**Fig. 2:**
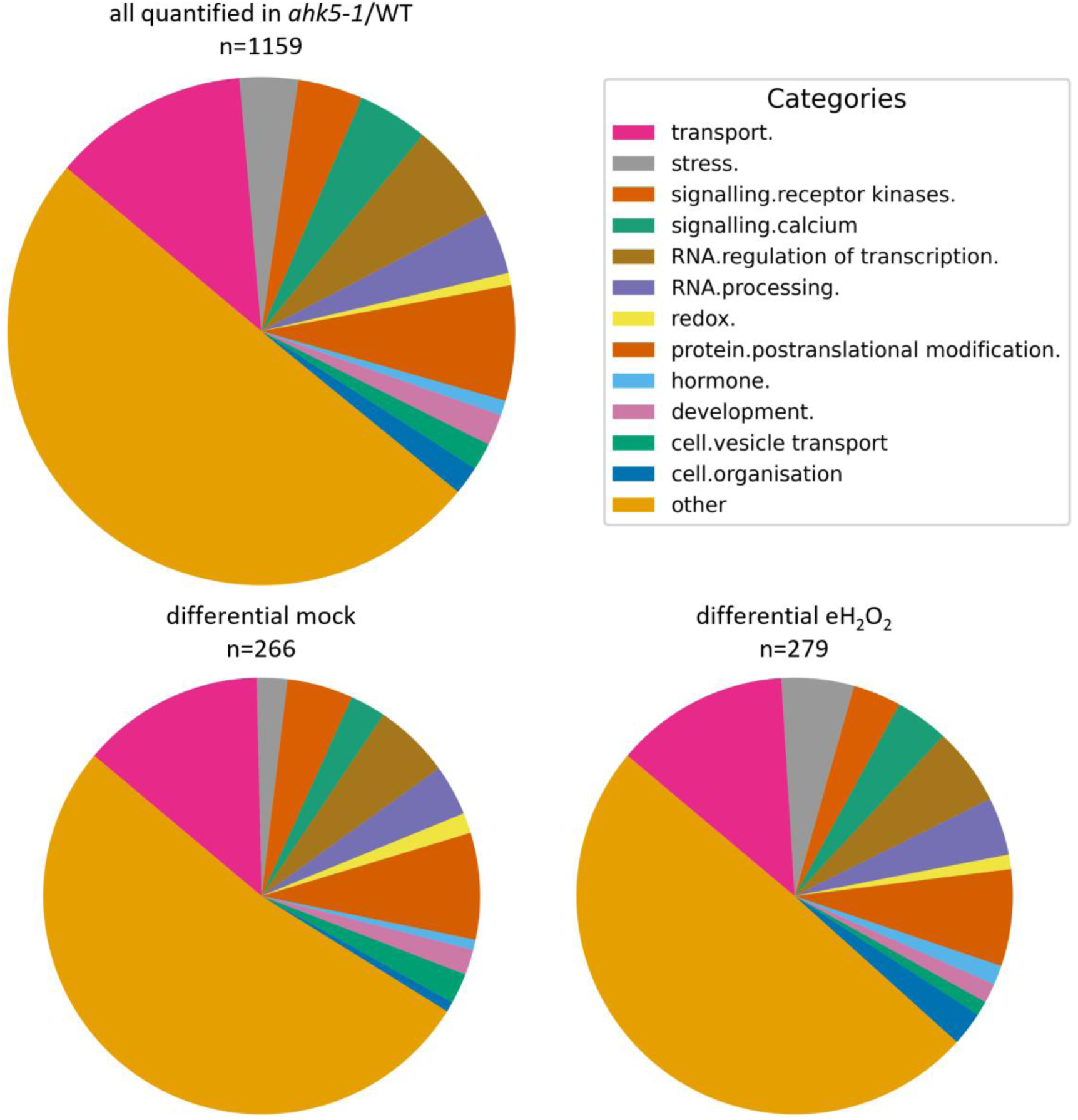
Functional categorization of all quantified phosphopeptides in *ahk5-1*. Quantified phosphopeptides categorized according to mapman in *ahk5-1* and differentially phosphorylated phosphopeptides in mock treated and eH2O2 treated *ahk5-1* samples. In total, 1159 phosphopeptides were quantified across all *ahk5-1* samples, of which 266 in mock experiments and 279 in eH2O2 experiments were rated differentially phosphorylated (|log2FC| >2) and categorized as indicated. The category “other” contains cell wall., cell.cycle, cell.division, DNA., gluconeogenesis / glyoxylate cycle., glycolysis., lipid.metabolism., major CHO metabolism., metal handling, minor CHO metabolism., misc., mitochondrial electron transport / ATP synthesis., N-metabolism., not assigned., nuceleotide.metabolism., protein.degradation., protein.folding, protein.glycosylation, protein.targeting., protein synthesis., PS., RNA.RNA binding, secondary metabolism., signalling.G-proteins, signalling.in sugar and nutrient physiology, signalling.light, signalling.MAP kinases, signalling.phosphinositides, and signalling.unspecified.

To gain a deeper understanding of AHK5-mediated responses to eH_2_O_2_ and to shed light on the potential biological relevance of quantified phosphopeptides, individual phosphorylation profiles were generated. Phosphopeptides were classified as “eH_2_O_2_ dependent mediated by AHK5”, if they met two conditions: A |log_2_FC| of phosphorylation levels of >2 between mutant and WT in either mock or eH_2_O_2_ treated experiments and a log_2_FC difference greater than 1 between mock and eH_2_O_2_ treated samples. Similarly, phosphopeptides were classified “eH_2_O_2_ independent mediated by AHK5”, if the log_2_FC difference between the two conditions was less than 1. Setting a threshold for phosphorylation level differences between eH_2_O_2_- and mock-treated phosphopeptides was done to confirm that the associated proteins are genuinely involved in the cellular response to eH_2_O_2_ (Fig. 3).

**Fig. 3:**
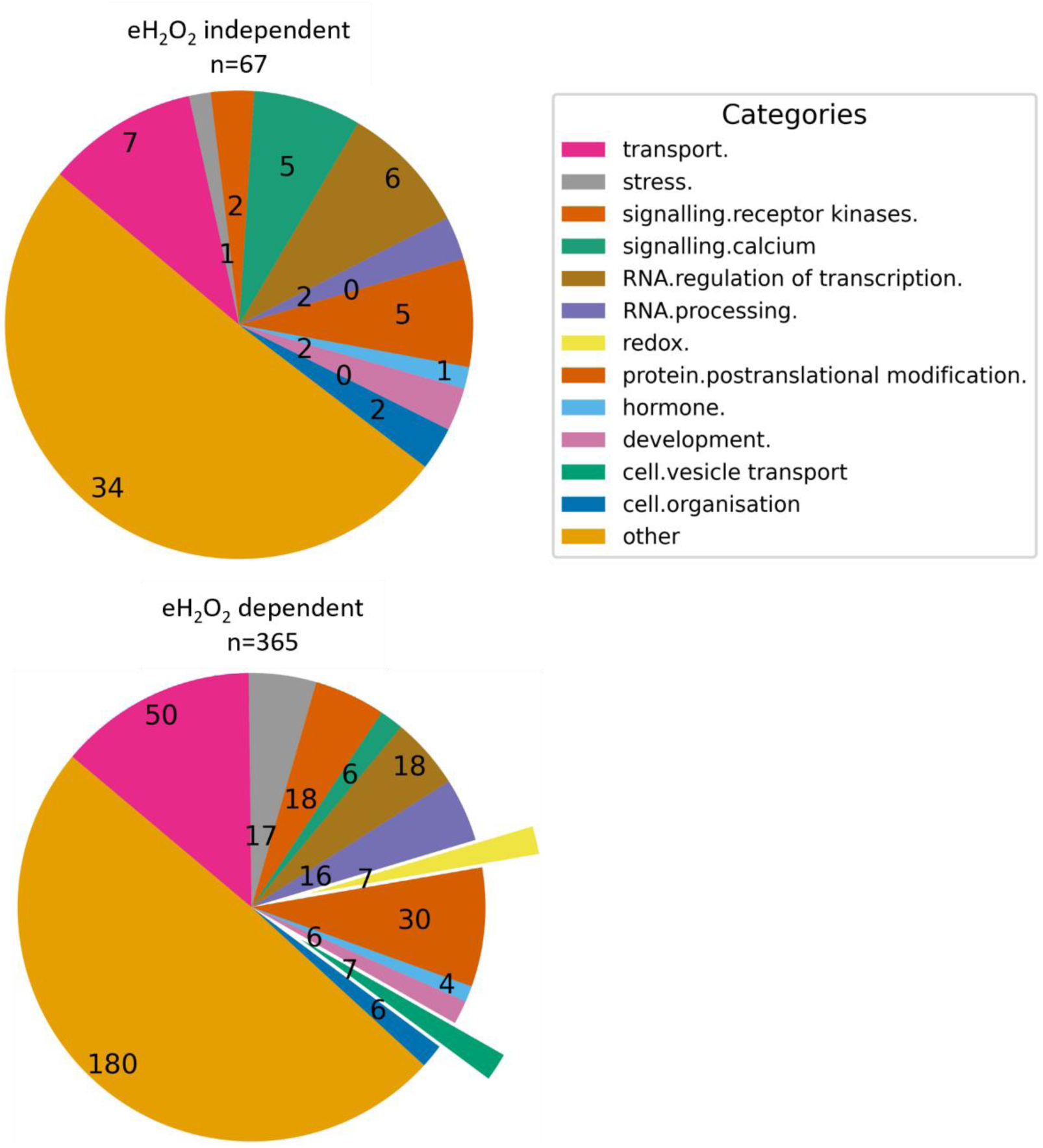
DPPs evaluated eH2O2 independent or dependent mediated by AHK5. 67 phosphopeptides were considered eH2O2 independent and 365 were considered eH2O2 dependent across all phosphopeptides, which were rated differentially phosphorylated (|log2FC| value >2) in the *ahk5-1* phosphoproteome in comparison to the WT phosphoproteome. Functional categorization as indicated. The category “other” contains the same sub-categories as Fig. 2.

Of all DPPs resulting from (altered) AHK5 function, comparably few were considered eH_2_O_2_ independent (n=67), whereas much more were considered eH_2_O_2_ dependent (n=365), (Fig. 3). The largest categories within the “eH_2_O_2_ dependent mediated by AHK5” phosphoproteins were assigned to “transport”, “RNA.”, “protein.postranslational modification.”, “signalling.receptor kinases”, and “stress”. Notably highlighted are the seven DPPs in the categories of “redox” and “cell/vesicle transport” that appeared exclusively in the “eH₂O₂-dependent” profiles and were not found in the “eH₂O₂-independent” profiles. Based on these phosphorylation profiles, the most interesting corresponding AHK5- and H_2_O_2_-regulated phosphoproteins are reviewed hereinafter, focusing on their biological functions.

### AHK5 likely modifies by phosphorylation the nanoscale organization of signaling hubs at the PM

As recently demonstrated by the identification of the extracellular H_2_O_2_ sensor HPCA1, H_2_O_2_ decoding can occur directly at the apoplast/PM interface. HPCA1 not only regulates stomatal aperture but also mediates cell-to-cell signaling in response to H_2_O_2_ (Wu *et al*., 2020; Fichman *et al*., 2022; Mishra *et al*., 2022). Emerging concepts suggest that ROS- (and Ca^2+^-) decoding nanodomains could facilitate localized signaling *via* the clustering of ROS-sensing and -decoding proteins at the cytosolic side of membranes. Signaling specificity and integration could be achieved through modulation of nanodomain organization with shared and overlapping core components of H_2_O_2_ and Ca^2+^ signaling systems (for reviews see (Demidchik and Shabala, 2018; Meyer *et al*., 2021; Wang *et al*., 2021); Hdedeh *et al*. (2025)).

Table 1 presents the phosphopeptides and associated proteins potentially linked to PM scaffold signalosomes differentially regulated by AHK5. Remorins, known to be primary nanodomain organizers at the PM, may play a key role in ROS signal integration, particular by their interaction with aquaporins which permeate H_2_O_2_ into the cytosol (Bienert *et al*., 2007; Dynowski *et al*., 2008; Demir *et al*., 2013; Fox *et al*., 2020; Hdedeh *et al*., 2025). The identification of several of these potential PM stress signalosome components to be differentially phosphorylated (Table 1) indicates, that AHK5 might play an important role in modulating these signalosomes and therefore the associated signaling pathways.

**Table 1:**
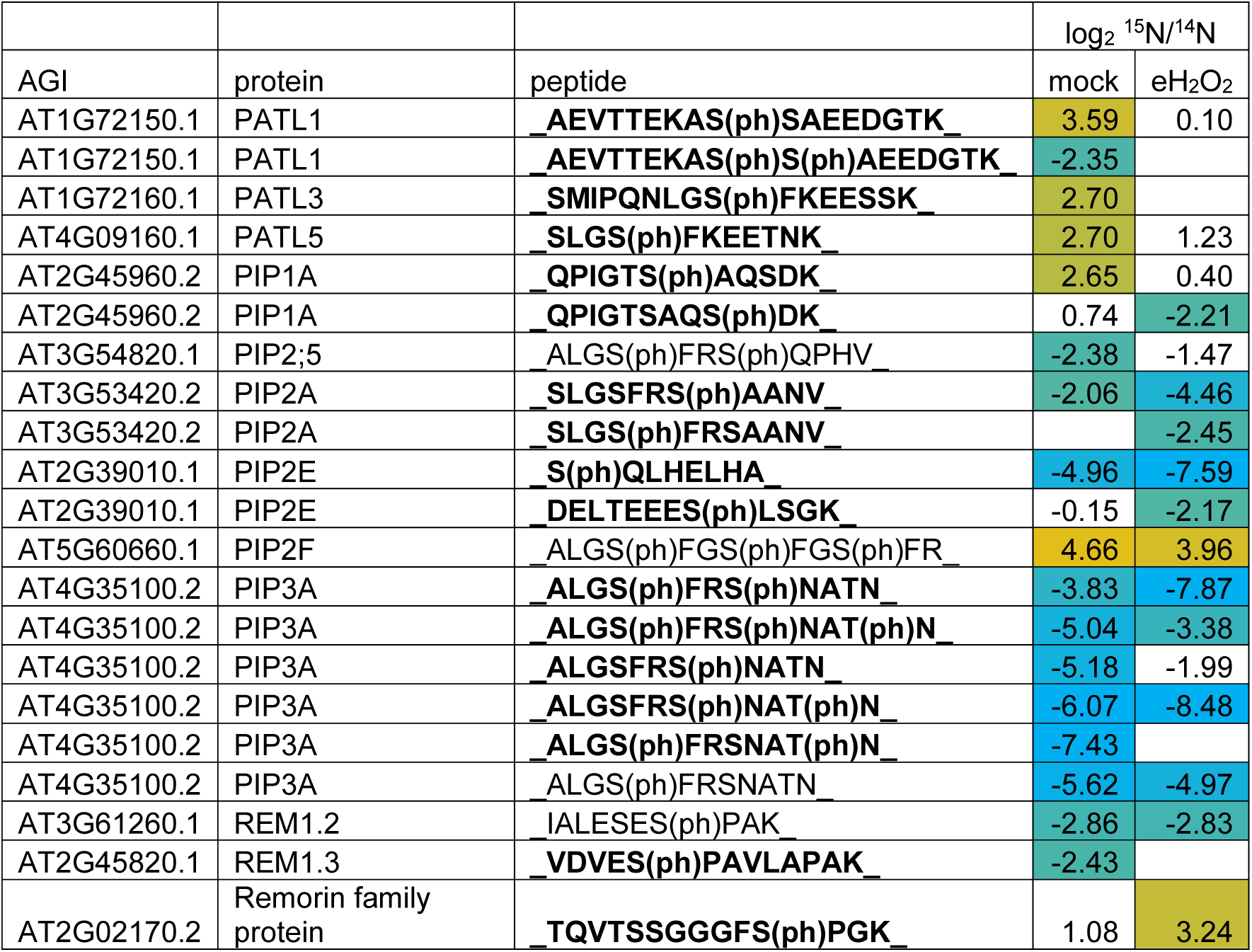
Selected DPPs in the *ahk5-1* phoshoproteome attributed to PM scaffolds. log2FC of interest of the *ahk5-1* phosphoproteome compared to WT phosphoproteome after 10 min of mock (mock) or 5 mM eH2O2 (eH2O2) treatment. Phosphorylated residues are indicated by a subsequent (ph), acetylation (ac) and oxidation (ox) are marked accordingly. Phosphopeptides in bold were assessed “eH2O2 dependent mediated by AHK5” and non-highlighted phosphopeptides were assessed “eH2O2 independent mediated by AHK5”. Absolute log2FC values above 2 are considered statistically significant and corresponding positive (orange) and negative (blue) values (i.e., respectively more or less phosphorylated, in *ahk5-1* compared to WT) are color graded limited at -6 and +6.

This hypothesis is further supported by the observation, that AHK5 exhibits nucleo-cytoplasmic localization with colocalization to the plasma membrane and is known to mediate various stress related responses. Furthermore, AHK5 contains a redox-active Cys3 residue that facilitates redox-dependent dimerization of its input domain *in vitro*, with a redox midpoint potential of -154 mV (Desikan *et al*., 2008; Pham and Desikan, 2012; Pham *et al*., 2012; Heunemann, 2016). This may enable AHK5 to rapidly detect transient fluctuations in the cytosolic redox potential in proximity to PM-associated signalosomes.

REM1.2 and REM1.3 and an uncharacterized REMORIN family protein (AT2G02170) exhibited AHK5-dependent phosphorylation (Table 1). Notably, phosphorylation of Ser13 of REM1.2 was not classified as H_2_O_2_-responsive in contrast to phosphorylation of Ser14 of REM1.3. REM1.3 was shown to interact with the RR ARR4 in yeast (Yamada *et al*., 1998), suggesting that AHK5 could affect the phosphostate of remorins through interaction with a RR through an unknown mechanism.

Furthermore, aquaporins, such as PIP1B, PIP2A, and PIP3A, were also identified as AHK5-regulated phosphoproteins. In PIP1B, Ser23 and Ser26, located in a cytosolic region, displayed eH_2_O_2_-dependent phosphorylation profiles. Similarly, PIP2A features two serines, Ser280 and Ser283, which are also located within a cytoplasmic domain at the C-terminus of PIP2A, that displayed AHK5- and eH_2_O_2_-dependent phosphorylation. Lower levels of phosphorylation of Ser283 in PIP2A in *ahk5-1* plants was even more pronounced following eH_2_O_2_ treatment (Table 1; Supplemental Fig. S1), suggesting that eH_2_O_2_ induced phosphorylation changes of aquaporins depend on the presence of functional AHK5 (Supplemental Fig. S1). In contrast to Ser280, phosphorylation of Ser283 increases in response to H_2_O_2_, but decreases in response to NaCl in WT plants and is crucial for PIP2A PM trafficking (Prak *et al*., 2008). This may indicate that AHK5’s negative regulation of salt tolerance (Pham *et al*., 2012) is caused by limiting dephosphorylation of Ser283 during salt stress.

Patellins, which were shown to be involved in plant membrane trafficking and cell wall formation affecting plant salt tolerance (Peterman *et al*., 2004; Peterman *et al*., 2006; Zhou *et al*., 2018; Sun *et al*., 2019; Zhou *et al*., 2019), were also among AHK5 dependent phosphoproteins. Specifically, PATL1 PATL3, and PATL5 featured several exclusively eH_2_O_2_-dependent phosphorylation sites (Table 1), implicating AHK5 function in the regulation of a PM hub that include remorins, aquaporins and patellins during stress responses.

Interestingly, both PIP1B and PIP2A were demonstrated in the past to interact with many AHK5-dependent phosphoproteins, including potential nanodomain organizers PATL1, REM1.2, and REM1.3 (Bellati *et al*., 2016). Collectively, these findings suggest that aquaporins, remorins, and patellins may represent core components of AHK5 regulated PM signalosomes, orchestrating stress responses through localized, dynamic signaling networks.

### Stomatal movement is likely influenced by AHK5-dependent protein phosphorylation

The most prominent phenotype observed in plants lacking functional AHK5 is the impaired stomatal closure in response to H_2_O_2_, ethylene (ET), darkness, and flg22 (Iwama *et al*., 2007; Desikan *et al*., 2008; Pham *et al*., 2012; Heunemann, 2016). In agreement with this phenotype, we identified several AHK5-dependent differentially phosphorylated proteins (DPs) and phosphopeptides (DPPs) associated with stomatal regulation (Table 2).

**Table 2:** Selected DPPs in the *ahk5-1* phoshoproteome attributed to stomatal regulation. log2FC of DPPs of interest of the *ahk5-1* phosphoproteome compared to WT phosphoproteome after 10 min of mock (mock) or 5 mM eH2O2 (eH2O2) treatment. Phosphorylated residues are indicated by a subsequent (ph), acetylation (ac) and oxidation (ox) are marked accordingly. Phosphopeptides in bold were assessed “eH2O2 dependent mediated by AHK5” and non-highlighted phosphopeptides were assessed “eH2O2 independent mediated by AHK5”. Absolute log2FC values above 2 are considered statistically significant and corresponding positive (orange) and negative (blue) values (i.e., respectively more or less phosphorylated, in *ahk5-1* compared to WT) are color graded limited at -6 and +6.

| AGI | protein | peptide | log<sub>2</sub> <sup>15</sup>N/<sup>14</sup>N |  |
| --- | --- | --- | --- | --- |
|  |  |  | mock | eH<sub>2</sub>O<sub>2</sub> |
| AT1G04120.2 | ABCC5 | **EVQEGGS(ph)ASDLK** | 0.14 | -2.06 |
| AT2G18960.1 | AHA1 | **T(ph)LHGLQPK** | -6.53 | -2.75 |
| AT5G23060.1 | CaS | **SFGTRS(ph)GT(ph)KFLPSSD** |  | -4.00 |
| AT4G23650.1 | CDPK6 (CPK3) | RGSSGS(ph)GTVGSSGSGTGGSR | -1.60 | -2.48 |
| AT4G04720.1 | CPK21 | **S(ph)GTITYEELK** |  | 2.18 |
| AT2G17290.1 | CPK6 | NSLNIS(ph)M(ox)R | 3.77 | 3.00 |
| AT5G03280.1 | EIN2 | **SLS(ph)GEGSGTGSLSR** | 2.71 |  |
| AT5G49760.1 | HPCA1 | **SLYDLQELLDTTIIASS(ph)GNLK** | 2.95 |  |
| AT5G49760.1 | HPCA1 | **SLYDLQELLDTTIIAS(ph)SGNLK** | 2.63 |  |
| AT5G49760.1 | HPCA1 | **S(ph)SIDAPQLMGAK** | -2.12 |  |
| AT5G12080.3 | MSL10 | S(ph)VGS(ph)PAPVTPSK | 2.02 | 1.47 |
| AT5G19520.1 | MSL9 | IPS(ph)PEGLVR | 2.54 | 3.29 |
| AT5G19520.1 | MSL9 | SVASAALS(ph)K | 1.64 | 2.45 |
| AT5G47910.1 | RBOHD | **ILSQMLS(ph)QK** | -1.36 | -2.45 |
| AT1G10940.1 | SNRK2.4 | **EVHAS(ph)GEVR** |  | -2.60 |

Stomatal closure involves sensing of extracellular Ca^2+^ and H_2_O_2_, triggering the intracellular increase of both second messengers and corresponding feedback mechanisms. Several proteins potentially involved in these processes, namely the potential mechanosensitive Ca^2+^ influx channels MSL9 and MSL10, and Ca^2+^ dependent kinases CPK3, 6, and 21, the NADPH oxidase RBOHD, the ABC transporter MRP5/ABCC5, and the calcium-sensing receptor CAS were identified as AHK5- and eH_2_O_2_-dependent phosphoproteins (Table 2) (Gaedeke *et al*., 2001; Mori *et al*., 2006; Suh *et al*., 2007; Ye *et al*., 2013; Chen *et al*., 2017; Yang *et al*., 2017). Here, not all proteins were eH_2_O_2_-dependent. For example, phosphorylation of CPK6, CDPK6 (CPK3), MSL9, and MSL10 were found to be up-regulated in the *ahk5-1* mutant independent of eH_2_O_2_ (Supplemental Fig. S2).

One of the key players of stomatal closure is RBOHD, which contributes to apoplastic H_2_O_2_ production. The AHK5-dependent phosphoresidue Ser347 of RBOHD is essential for its activity and is also known to be phosphorylated by BIK1 and CPKs (Kadota *et al*., 2014; Lee *et al*., 2020). In *ahk5-1* plants, phosphorylation at this residue was reduced, particularly following eH_2_O_2_ treatment (Table 2), suggesting that AHK5 may enhance RBOHD activity, in response to eH_2_O_2_. Notably, additional phosphorylation sites of RBOHD known responsive to pathogen and wounding associated signals like flg22 and ATP (Nühse *et al*., 2007; Dubiella *et al*., 2013; Lee *et al*., 2020), were not differentially regulated, supporting an AHK5 mediated pathway specifically activated by eH_2_O_2_. In the established ABA-dependent stomatal closure pathway, RBOHD is probably phosphorylated by OST1 at position 347, which activates anion channels SLAC1 and QUAC1 (Geiger *et al*., 2009; Lee *et al*., 2009; Imes *et al*., 2013; Fichman *et al*., 2022). However, these proteins were not differentially phosphorylated in *ahk5-1* plants compared to WT, fitting to the AHK5-dependent pathway operating independently of ABA (Desikan *et al*., 2008). Instead, OST1-LIKE 7 (OSKL7/SnRK2.4) was identified as potential AHK5 phosphorylation target (Table 2), suggesting an alternative signaling route. Consistently, SLAC1-HOMOLOG PROTEIN 3 (SLAH3), an anion efflux channel, showed reduced phosphorylation of Ser128 and Ser601 in *ahk5-1* under mock conditions compared to WT, though changes were below the significance threshold chosen (Supplemental Table S1). Both sites were shown to inhibit SLAH3 function when phosphorylated (Liu *et al*., 2019; Sun *et al*., 2021), implying AHK5-promoted release of SLAH3 inhibition after H_2_O_2_ perception. Importantly, SLAH3 has been shown to function downstream of CPK3, CPK6, and CPK21, the only CPKs demonstrated to phosphorylate both SLAC1 and SLAH3 (Geiger *et al*., 2010; Brandt *et al*., 2012; Scherzer *et al*., 2012; Kollist *et al*., 2014). Thus, a potential AHK5-SnRK2.4-CPK-SLAH3 module may represent a novel signaling pathway that parallels the classical ABA-OST1-SLAC1 module.

CAS has been proposed to initiate chloroplast-derived H_2_O_2_ production to drive stomatal closure in response to elevated extracellular Ca^2+^ (Nomura *et al*., 2008; Weinl *et al*., 2008; Wang *et al*., 2011). Here, CAS displayed AHK5-promoted phosphorylation at two serines specifically upon eH_2_O_2_ treatment (Table 2), suggesting its participation in the AHK5-mediated stomatal closure pathway.

Together, a possible signaling system emerges: extracellular H_2_O_2_ enters the cytosol via aquaporins, activating AHK5 once a critical ROS threshold is reached. AHK5 then enhances RBOHD activity via Ser347 phosphorylation, amplifying H_2_O_2_ production.

This promotes anion efflux via de-repressed SLAH3, triggering the activation of Ca^2+^ channels and further Ca²⁺ influx. The rise in cytosolic Ca²⁺ subsequently activates CAS, reinforcing chloroplast H_2_O_2_ production culminating in stomatal closure. This AHK5-dependent pathway would bypass ABA to initiate stomatal closure.

HPCA1 was recently identified as a key component required for eH_2_O_2_-induced cytosolic Ca^2+^ elevation and stomatal closure (Wu *et al*., 2020). In *ahk5-1*, three phosphosites in HPCA1 were differentially phosphorylated under non-stress conditions (Table 2). Among these, Ser606 was previously shown to be H_2_O_2_-responsive (Wu *et al*., 2020) and showed increased phosphorylation in presence of AHK5 under mock conditions. The other two sites, Ser866 and Ser867, located within the predicted kinase domain, represent novel phosphosites. Intriguingly, the activity of HPCAL1 (At5G49770), a root-expressed homolog of HPCA1, was shown to be enhanced when phosphorylated at the homologous Ser870 (Li *et al*., 2024). These findings suggest that AHK5 may modulate HPCA1 through a yet undefined mechanism under non-stress conditions, that is disrupted in response to eH_2_O_2_. Notably, recent findings propose that HPCA1 might be dispensable for the initiation of ROS signaling, but crucial for its systemic intercellular propagation. The functional relevance of HPCA1 regulation by AHK5 and the role of Ser866 and Ser867 phosphorylation, remains to be elucidated.

### AHK5 fine-tunes root growth and development by differential protein phosphorylation

Root development is regulated by a complex interplay of abscisic acid (ABA), ethylene (ET), and brassinosteroid (BR) signaling pathways. These pathways are closely linked to ROS homeostasis and Ca^2+^ influx, which together drive root elongation and establish root hair initiation sites (Foreman *et al*., 2003; Demidchik *et al*., 2007; Takeda *et al*., 2008; Hernández-Barrera *et al*., 2015; Lv *et al*., 2018).In addition to impaired stomatal regulation, *ahk5* mutants also exhibit defective root growth, which, unlike AHK5-mediated stomatal closure, is dependent on ABA signaling (Iwama *et al*., 2007).

AHK5 appears to modulate ET signaling via the ETR1-EIN2 pathway, which is also involved in root growth and development. Notably, *ein2* mutants show longer roots under non-stress conditions (Wang *et al*., 2007), while *ahk5* mutants are hypersensitive to ABA and, to a lesser extent, to the ET precursor ACC (Iwama *et al*., 2007). Ser650 of the central ET signaling regulator ETHYLENE-INSENSITIVE PROTEIN 2 (EIN2) exhibited AHK5- and H_2_O_2_-dependent phosphorylation (Table 3). EIN2 plays a pivotal role in a wide range of ET-mediated response, including stomatal regulation, senescence, root development, and defense responses (Bouchez *et al*., 2007; Wang *et al*., 2007; Wang *et al*., 2020; Yu *et al*., 2021; Zhang *et al*., 2021). EIN2 activity is repressed by ETR1, which promotes interaction with CTR1 in its phosphorylated state, thereby preventing EIN2 activation leading to its degradation (Bisson *et al*., 2009; Bisson and Groth, 2010). Thus, AHK5-mediated root development likely relies on both ET and ABA signaling, suggesting a redox-sensing mechanism involving AHK5-triggered Ca^2+^ influx in a positive feedback loop with AHK5-dependent differential phosphorylation of aquaporins, remorins, different CPKs, and SLAH3, as discussed earlier, all of which play a role in root development as well (Boursiac *et al*., 2005; Zheng *et al*., 2015; Cubero-Font *et al*., 2016; Zou *et al*., 2016; Ke *et al*., 2021).

**Table 3:**
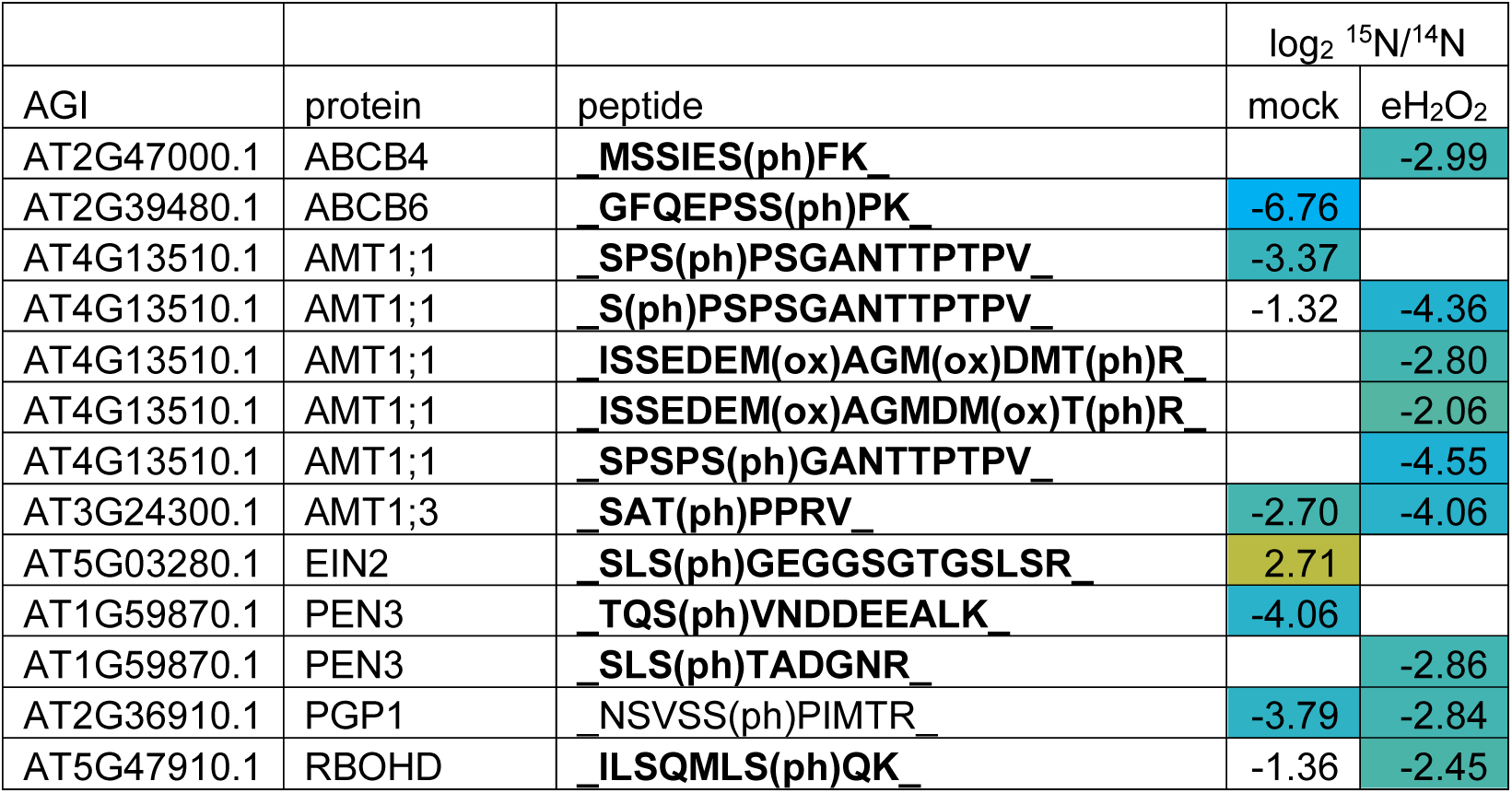
Selected DPPs in the *ahk5-1* phoshoproteome attributed to root growth. log2FC of DPPs of interest of the *ahk5-1* phosphoproteome compared to WT phosphoproteome after 10 min of mock (mock) or 5 mM eH2O2 (eH2O2) treatment. Phosphorylated residues are indicated by a subsequent (ph), acetylation (ac) and oxidation (ox) are marked accordingly. Phosphopeptides in bold were assessed “eH2O2 dependent mediated by AHK5” and non-highlighted phosphopeptides were assessed “eH2O2 independent mediated by AHK5”. Absolute log2FC values above 2 are considered statistically significant and corresponding positive (orange) and negative (blue) values (i.e., respectively more or less phosphorylated, in *ahk5-1* compared to WT) are color graded limited at -6 and +6.

| AGI | protein | peptide | $\log_2^{15}\text{N}/^{14}\text{N}$ | |
| --- | --- | --- | --- | --- |
| | | | mock | $\text{eH}_2\text{O}_2$ |
| AT2G47000.1 | ABCB4 | **MSSIES(ph)FK** |  | -2.99 |
| AT2G39480.1 | ABCB6 | **GFQEPSS(ph)PK** | -6.76 |  |
| AT4G13510.1 | AMT1;1 | **SPS(ph)PSGANTTPTPV** | -3.37 |  |
| AT4G13510.1 | AMT1;1 | **S(ph)PSPSGANTTPTPV** | -1.32 | -4.36 |
| AT4G13510.1 | AMT1;1 | **ISSEDEM(ox)AGM(ox)DMT(ph)R** |  | -2.80 |
| AT4G13510.1 | AMT1;1 | **ISSEDEM(ox)AGMDM(ox)T(ph)R** |  | -2.06 |
| AT4G13510.1 | AMT1;1 | **SPSPS(ph)GANTTPTPV** |  | -4.55 |
| AT3G24300.1 | AMT1;3 | **SAT(ph)PPRV** | -2.70 | -4.06 |
| AT5G03280.1 | EIN2 | **SLS(ph)GEGSGTGSLSR** | 2.71 |  |
| AT1G59870.1 | PEN3 | **TQS(ph)VNDDEEALK** | -4.06 |  |
| AT1G59870.1 | PEN3 | **SLS(ph)TADGNR** |  | -2.86 |
| AT2G36910.1 | PGP1 | **NSVSS(ph)PIMTR** | -3.79 | -2.84 |
| AT5G47910.1 | RBOHD | **ILSQMLS(ph)QK** | -1.36 | -2.45 |

NADPH oxidases like RBOHC, RBOHD, and RBOHF are critical in regulating root development by modulating ROS levels, with some evidence placing ROS upstream of Ca^2+^ influx (Foreman *et al*., 2003; Kwak *et al*., 2003; Takeda *et al*., 2008). Specifically, RBOHD and RBOHF negatively regulate lateral root formation independently of auxin (Li *et al*., 2015), while ABA-induced ROS through RBOHD and RBOHF inhibits primary root growth and duces AUX sensitivity, guiding root branching and root hair initiation (Jiao *et al*., 2013; Huang *et al*., 2020). Wound-induced ROS has also been shown to promote adventitious root formation by altering AUX synthesis and transport, likely involving AHK5-RBOH interactions. Several AUX-related proteins were identified as AHK5-dependent phosphoproteins, including ABC-type AUX efflux transporters such as PGP1. Phosphorylation of Ser634 of PGP1 was significantly reduced in *ahk5-1* under mock and eH_2_O_2_ conditions Table 3. Since phosphomimic and phosphodead mutations at this site increase and reduce AUX efflux, respectively (Henrichs *et al*., 2012), it is likely that AHK5 promotes AUX transport. This is substantiated by additional AUX transporters like ABCB4, ABCB6, and the AUX precursor transporter PEN3, which also displayed AHK5- and eH_2_O_2_-dependent phosphorylation profiles (Table 3).

Lateral root branching is promoted by ammonium availability and dependent on AMT1;3 (Lima *et al*., 2010). Thr494 of AMT1;3 was suggested as a phosphoresidue involved in the sophisticated fine-tuning and balancing of ammonium uptake, moderately reducing AMT1;3 facilitated NH^4+^ uptake (Wu *et al*., 2019). Thr494 showed AHK5 promoted phosphorylation with a more pronounced effect after eH_2_O_2_ treatment ^T^able 3. Strikingly, a second threonine in AMT1;3 at position 464 (Thr464) that completely inhibits AMT1;3 activity upon phosphorylation did not show an AHK5-dependent phosphorylation profile, indicating that AHK5 functions in fine-tuning, rather than full inhibition, of lateral root development and ammonium uptake. We also observed a consistent down-regulation of the inhibitory phosphorylation site at T460 in the *ahk5-1* mutant, and other regulatory phosphorylation sites in the C-terminus of AMTs, which were partly enhanced by eH_2_O_2_ (Supplemental Fig. S3). Nitrogen uptake has previously been linked to H_2_O_2_ signaling, for example through the activating phosphorylation of NRT2.1 by the H_2_O_2_ sensor HPCAL1 (Li *et al*., 2024). Interestingly, AMT1;3 interacts with numerous proteins identified as AHK5-dependent phosphoproteins, most notably PIP1B, PIP2A, and AMT1;1 (Bellati *et al*., 2016). AMT1;1 in turn also interacts with, PIP1B, and PIP2A, (Yuan *et al*., 2013; Jones *et al*., 2014; Bellati *et al*., 2016). PIP1B and PIP2A in turn interact with many AHK5-dependent phosphoproteins, such as CPKs, remorins, and patellins. These aquaporins may represent centrals nodes of AHK5 function, potentially anchoring signaling scaffolds in conjunction with remorins to modulate the activity associated PM signalosomes in response to H_2_O_2_ during root development. However, the dependency of AHK5-mediated root development on ABA indicates that some components may differ compared to AHK5-driven stomatal responses, e.g. the lack of CAS expression in roots as a potential ABA bypass to trigger high-level ROS production.

### Modifying activity of AHK5 on cytoskeleton scaffolds and trafficking

Stress responses, both biotic and abiotic, involve substantial alterations in cellular metabolism and composition. Central to the orchestration of adaptive responses is the dynamic reorganization of cytoskeletal scaffolds and intracellular trafficking, which modulate the assembly of PM-associated signalosomes (for review see Hdedeh *et al*. (2025)). This remodeling is accomplished by clathrin-mediated endocytosis (CME), which facilitates vesicle formation and internalization of cargo proteins in nanodomains to modulate PM signaling (Johnson and Vert, 2017; Kaksonen and Roux, 2018).

The phosphorylation of dynamins was found to be lower in the *ahk5-1* mutant compared to WT, while phosphorylation of cellulose synthase and PBL1 was upregulated in the mutant compared to WT (Table 4, Supplemental Fig. S3). Two dynamin related proteins, DRP2A (ADL6) and DRP2B (ADL3), have been shown to participate in CME, localizing to sterol-enriched membrane domains at root hair initiation sites (Fujimoto *et al*., 2010; Stanislas *et al*., 2015). Both ADL3 and ADL6 interact with PIP1B and PIP2A (Bellati *et al*., 2016), and were identified as AHK5-dependent phosphoproteins (Table 4), suggesting that AHK5 contributes not only to the modulation of nanodomain downstream signaling, but also the composition thereof. Additionally, cellulose synthase complexes (CESAs) are known to play a critical role in nanodomain organization as well. AHK5-regulated phosphoproteins such as RBOHD, AMT1;3, and PIP2A colocalize with CME components forming endocytic ‘hotspots’, that could facilitate stress signal integration and resulting protein turnover (for review see Hdedeh *et al*. (2025)). Several additional phosphoproteins involved in membrane tethering were identified to be modified in an AHK5-dependent manner, including IQD2 and IQD14 (Table 4). IQDs have been linked to both Ca^2+^ and AUX signaling to regulate the microtubule skeleton and membrane domain organization, thus influencing e.g. salt tolerance (Bürstenbinder *et al*., 2017a; Bürstenbinder *et al*., 2017b; Wendrich *et al*., 2018 [Preprint]; Yang *et al*., 2018 [Preprint]).

**Table 4:** Selected DPPs in the *ahk5-1* phoshoproteome attributed to cytoskeleton and trafficking. log2FC of DPPs of interest of the *ahk5-1* phosphoproteome compared to WT phosphoproteome after 10 min of mock (mock) or 5 mM eH2O2 (eH2O2) treatment. Phosphorylated residues are indicated by a subsequent (ph), acetylation (ac) and oxidation (ox) are marked accordingly. Phosphopeptides in bold were assessed “eH2O2 dependent mediated by AHK5” and non-highlighted phosphopeptides were assessed “eH2O2 independent mediated by AHK5”. Absolute log2FC values above 2 are considered statistically significant and corresponding positive (orange) and negative (blue) values (i.e., respectively more or less phosphorylated, in *ahk5-1* compared to WT) are color graded limited at -6 and +6.

| AGI | protein | peptide | log<sub>2</sub> <sup>15</sup>N/<sup>14</sup>N |  |
| --- | --- | --- | --- | --- |
|  |  |  | mock | eH<sub>2</sub>O<sub>2</sub> |
| AT1G59610.1 | ADL3 | **AAAASSWSDNSGTES(ph)SPR** |  | -5.72 |
| AT1G59610.1 | ADL3 | **AAAASSWSDNSGTESS(ph)PR** |  | -4.45 |
| AT1G59610.1 | ADL3 | **QLS(ph)IHDNR** |  | -3.35 |
| AT1G10290.1 | ADL6 | **GQDAEQSLLS(ph)R** | 4.76 |  |
| AT1G10290.1 | ADL6 | **AT(ph)SPQPDGPTAGGSLK** | 3.92 | 0.58 |
| AT1G10290.1 | ADL6 | **ATS(ph)PQPDGPTAGGSLK** | -2.67 | -4.39 |
| AT1G10290.1 | ADL6 | **AAAASSYSDNSGTES(ph)SPR** | -1.63 | -3.37 |
| AT5G05170.1 | CESA3 | LPYSSDVNQS(ph)PNR | 5.27 | 4.29 |
| AT5G05170.1 | CESA3 | **NTGPVSTQAAS(ph)ER** | -2.54 |  |
| AT5G05170.1 | CESA3 | **NTGPVS(ph)TQAASER** | -0.49 | 4.63 |
| AT2G43680.3 | IQD14 | **GPGS(ph)PGGVVLEK** | 1.93 | 3.92 |
| AT2G43680.3 | IQD14 | TLS(ph)PKPPS(ph)PR | -3.10 | -2.37 |
| AT5G03040.3 | iqd2 | QSSS(ph)SPPPALAPR | -2.86 | -2.87 |
| AT3G55450.1 | PBL1 | **SSS(ph)GLVIAVK** | 2.43 |  |
| AT3G55450.1 | PBL1 | **TEGEILS(ph)STTVK** |  | 2.79 |
| AT5G56460.1 | PBL16 | **SES(ph)PKEQSPTVEDK** | 2.62 |  |
| AT5G47070.1 | PBL19 | **SETS(ph)SFNLQTPR** |  | 4.14 |
| AT5G47070.1 | PBL19 | **SETSS(ph)FNLQTPR** |  | 2.55 |
| AT1G78880.1 | SHOU4 | **TNS(ph)GPLSK** | 2.11 |  |
| AT1G78880.1 | SHOU4 | **MS(ph)GNLASAGS(ph)NSMKK** | 1.07 | 2.65 |

PATL1 and PATL3, as described before, featured newly discovered AHK5-dependent phosphosites as shown in Table 1. PATL1 is involved in membrane trafficking at the cell plate during cytokinesis and cell wall formation (Peterman *et al*., 2004), while PATL3 has been implicated in virus defense (Peiro *et al*., 2014). Collectively, patellins participate in development and stress responses, possibly by modulating AUX fluxes through PIN relocalization (Tejos *et al*., 2018; Zhou *et al*., 2019). However, no PIN proteins were found to be AHK5- and eH_2_O_2_-dependent phosphoproteins. In contrast, several ABC-type transporters were AHK5- and eH_2_O_2_-dependent phosphoproteins, suggesting that patellins may influence AHK5-mediated auxin transport during root development under stress conditions via ABC transporters instead. Moreover, CESA3, which functions in cell wall and stress signaling likely mediated by jasmonate and ethylene (Ellis *et al*., 2002; Caño-Delgado *et al*., 2003), was found to contain three AHK5-regulated phosphoserines: Ser167, Ser211, and Ser 216 (Table 4). While Ser167 did not show response to eH_2_O_2_ treatment, Ser211 and Ser216 did. Interestingly, SHOU4, a novel PM-associated protein, implicated in CESA trafficking during endocytosis (Polko *et al*., 2018) was also differentially phosphorylated in an AHK5-dependent manner at two sites (Table 4) with several additional sites showing changes that did not meet the chosen threshold for significance.

Collectively, these findings point to a potential AHK5-regulated pathway modulating CESA trafficking via SHOU4 and SECs along the microtubule cytoskeleton. Exocytic vesicles carrying CESAs may be targeted to specific PM regions by scaffolding proteins such as PATELLINs, REMORINs, and IQDs. This targeted delivery likely contributes to the modulation of cell wall architecture, CME, and thus nanodomain signalosomes in response to H_2_O_2_, produced in response to stress or developmental cues.

Three PBS1-LIKE PROTEINS (PBLs) PBL1, PBL16, and PBL19 were also found to be AHK5- and eH_2_O_2_-dependent phosphoproteins (Table 4). PBL1, which shares functional redundancy with BIK1, regulates callose deposition and inhibits root growth in response to flg22 (Zhang et al., 2010; Ranf et al., 2014). Given PBL1’s role in defense signaling and the involvement of BIK1 in salicylic acid (SA) regulation (Veronese et al., 2005; Lal et al., 2018), it is plausible that AHK5 contributes to SA-mediated signaling and systemic acquired resistance (SAR). The identification of PBL1, PBL16, and PBL19 as AHK5 targets suggests a regulatory role for AHK5 in the phosphorylation of RBOHD and potentially other immune signaling components.

### Conclusions and perspectives

In essence, we revealed a massive signaling transition from the AHK5-dependent TCS phosphorelay to Ser/Thr phosphorylation/dephosphorylation. Against the background that it may function as an intracellular redox sensor (Drechsler, 2022; Nöldeke, 2022), AHK5 fine-tunes many aspects of H_2_O_2_-, redox potential- and Ca^2+^-dependent stress decoding processes by posttranslational modification. Supported by published protein-protein interaction data and the fast modification of for instance PM-associated PIPs, PATLs, REMs, IQDs, aquaporins (PIPs), various transporters and other proteins, reported here, AHK5 may be a constituent of a stress signaling and integrating hub (signalosome) at the PM/cytosol interface that balances biotic and abiotic stress responses against internal processes such as root growth and development as well as stomatal closure. Regarding the establishment of specificity in TCS-regulated signaling, it is of great interest that the global HK-controlled phosphorylation pattern differs between AHK1, AHK3/4 and AHK5 (Dautel, 2016; Dautel *et al*., 2016). This suggests, that different HK-dependent TCS processes are structurally intertwined with other phosphorylation-controlled signaling networks in distinct signalosomes.

In future, it will be crucial exploring (1) how and at which level within the AHK5-controlled TCS (HK, HP and/or RR) the signaling transition occurs, (2) what the consequences of the newly identified modifications are for the functional properties of the targeted proteins, and (3) how the entire phosphorelay-phosphorylation network is organized regarding the proposed signalosome concept.

## Supporting information

List of phosphopeptides and comparisons

## Supplementary data

Table S1: Complete list of all quantified phosphopeptides in the *ahk5-1* mutant and wild type (WT).

Figure S1: Phopshopeptides matching scaffold proteins and aquaporins highlighted in the cross plot of untreated and H_2_O_2_ treated plants.

Figure S2: Phopshopeptides matching proteins attributed to stomatal regulation highlighted in the cross plot of untreated and H_2_O_2_ treated plants.

Figure S3: Phopshopeptides matching proteins attributed to root growth and cellular trafficking highlighted in the cross plot of untreated and H_2_O_2_ treated plants.

## Acknowledgements

We thank Radhika Desikan for providing *ahk5-1* seeds.

## Author contributions

TD and KH designed the research and experiments. TD, ZL and WXS performed the experiments and data analyses. TD wrote the manuscript, and WXS as well as KH proofread the manuscript.

## Data availability

All data supporting the findings are available in the supplementary data and from the corresponding author.

---

**Fig. S1:**
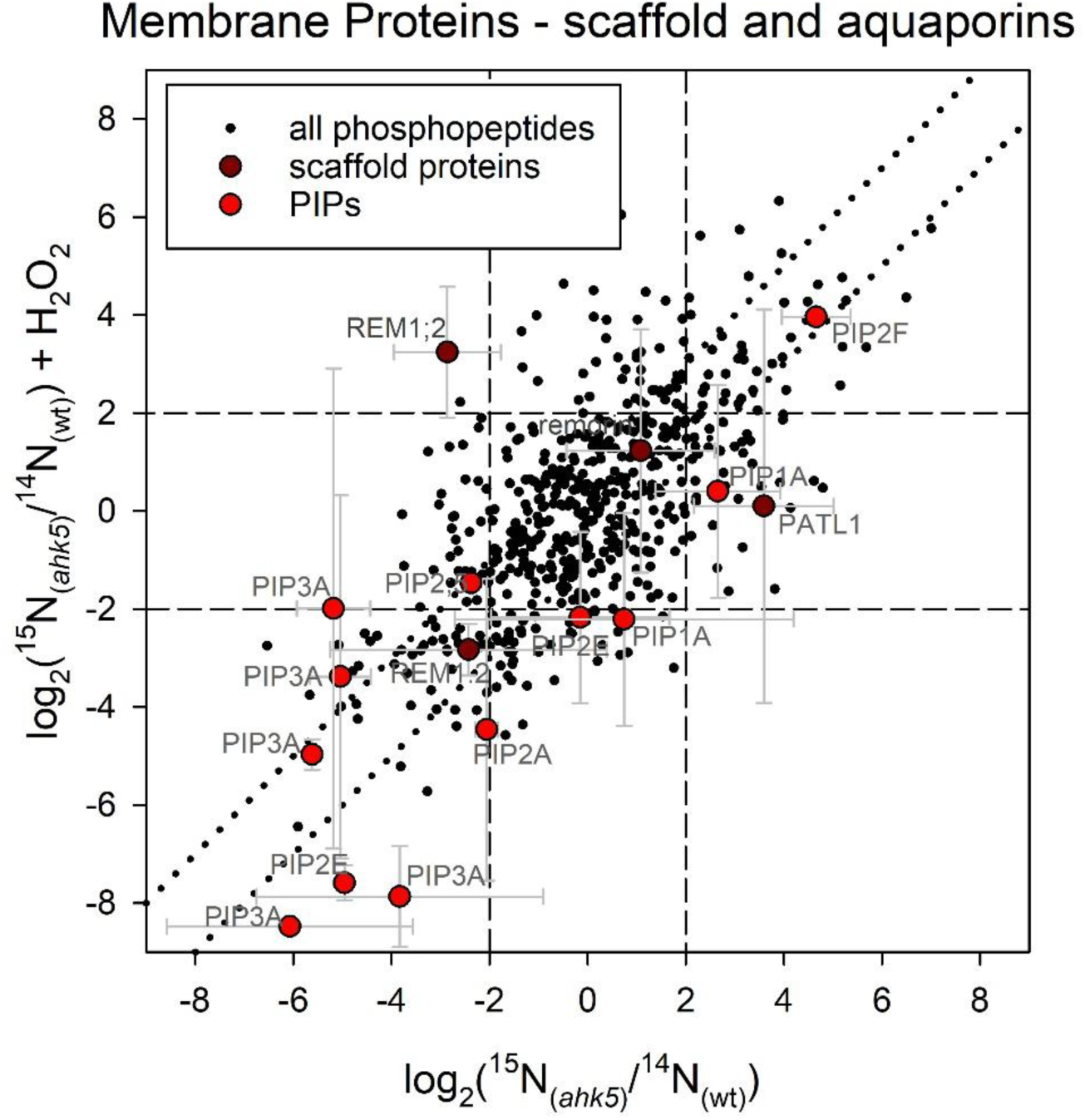
Phopshopeptides matching scaffold proteins and aquaporins in the cross plot of untreated and H2O2 treated plants. Details see tables.

**Fig. S2:**
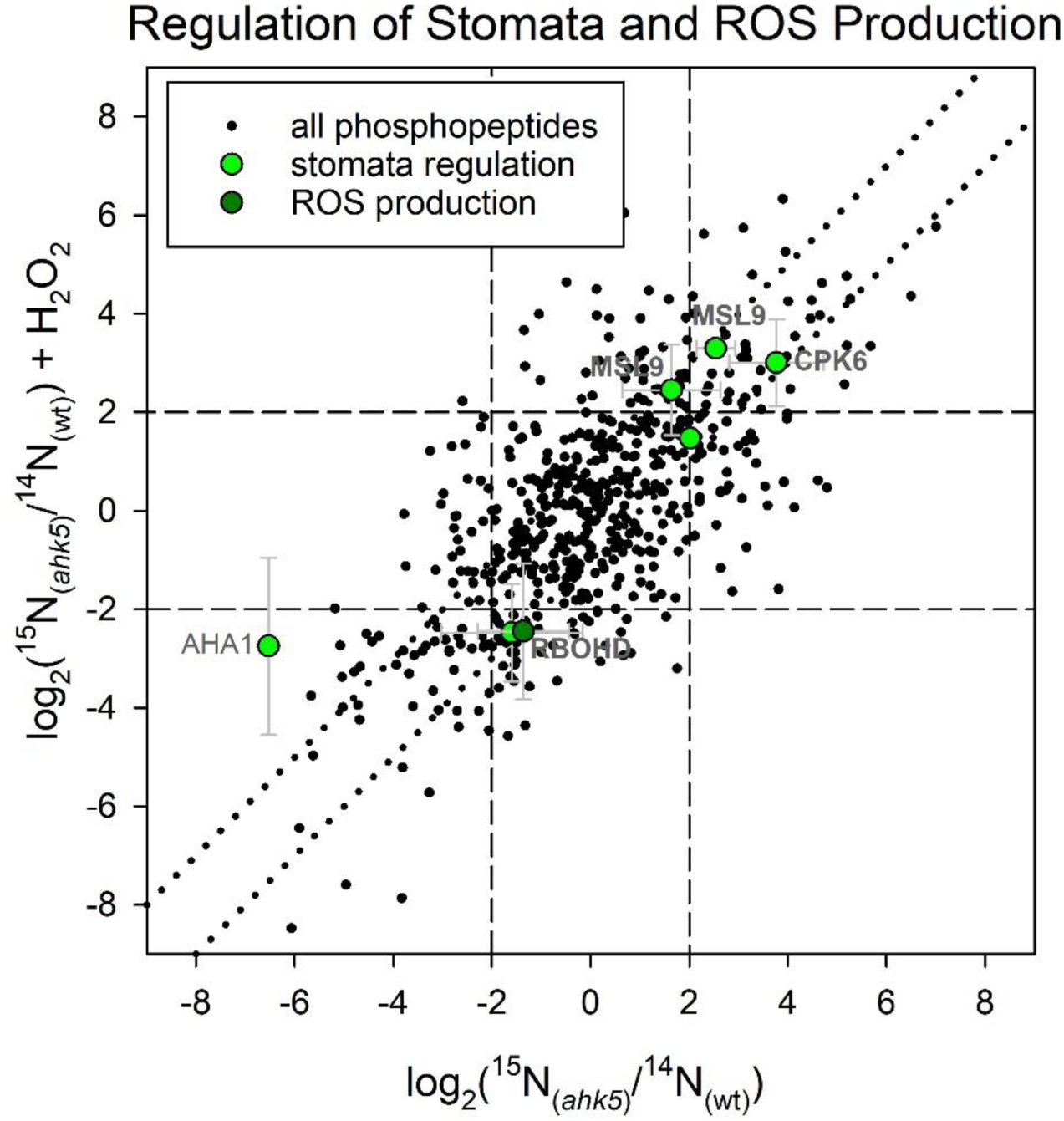
Phopshopeptides matching proteins attributed to stomatal regulation in the cross plot of untreated and H2O2 treated plants. Details see tables.

**Fig. S3.**
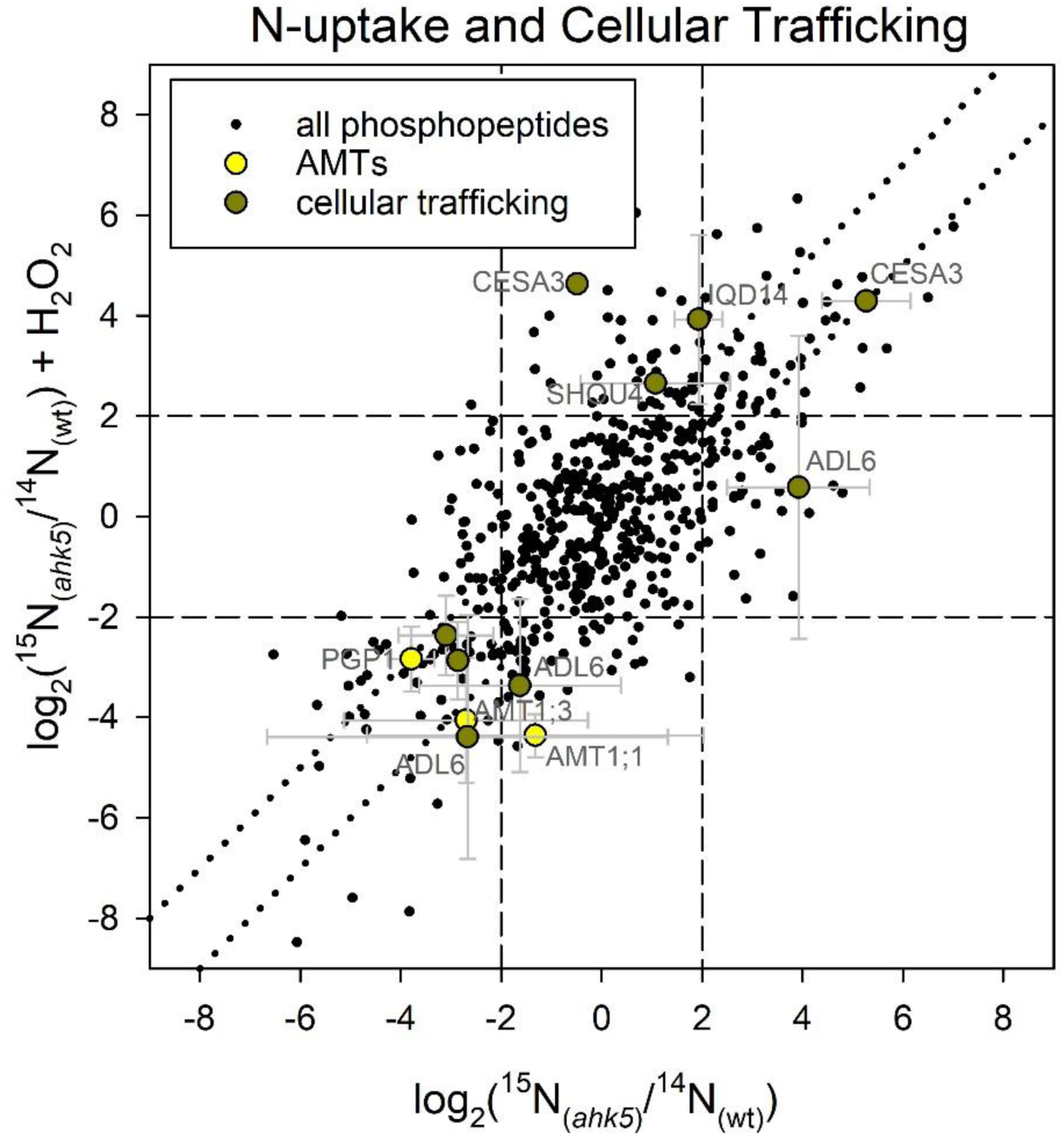
Phopshopeptides matching proteins attributed to N-uptake and cellular trafficking in the cross plot of untreated and H2O2 treated plants. Details see tables.

